# Sensitive detection of synthetic response to cancer immunotherapy driven by gene paralog pairs

**DOI:** 10.1101/2024.07.02.601809

**Authors:** Chuanpeng Dong, Feifei Zhang, Emily He, Ping Ren, Nipun Verma, Xinxin Zhu, Di Feng, Hongyu Zhao, Sidi Chen

## Abstract

Emerging immunotherapies such as immune checkpoint blockade (ICB) and chimeric antigen receptor T-cell (CAR-T) therapy have revolutionized cancer treatment and have improved the survival of patients with multiple cancer types. Despite this success many patients are unresponsive to these treatments or relapse following treatment. CRISPR activation and knockout (KO) screens have been used to identify novel single gene targets that can enhance effector T cell function and promote immune cell targeting and eradication of tumors. However, cancer cells often employ multiple genes to promote an immunosuppressive pathway and thus modulating individual genes often has a limited effect. Paralogs are genes that originate from common ancestors and retain similar functions. They often have complex effects on a particular phenotype depending on factors like gene family similarity, each individual gene’s expression and the physiological or pathological context. Some paralogs exhibit synthetic lethal interactions in cancer cell survival; however, a thorough investigation of paralog pairs that could enhance the efficacy of cancer immunotherapy is lacking. Here we introduce a sensitive computational approach that uses sgRNA sets enrichment analysis to identify cancer-intrinsic paralog pairs which have the potential to synergistically enhance T cell-mediated tumor destruction. We have further developed an ensemble learning model that uses an XGBoost classifier and incorporates features such as gene characteristics, sequence and structural similarities, protein-protein interaction (PPI) networks, and gene coevolution data to predict paralog pairs that are likely to enhance immunotherapy efficacy. We experimentally validated the functional significance of these predicted paralog pairs using double knockout (DKO) of identified paralog gene pairs as compared to single gene knockouts (SKOs). These data and analyses collectively provide a sensitive approach to identify previously undetected paralog pairs that can enhance cancer immunotherapy even when individual genes within the pair has a limited effect.

## Introduction

Emerging immunotherapies, especially immune checkpoint blockade (ICB) and adoptive cell therapy, have revolutionized cancer treatment for multiple cancer types; however, a significant fraction of patients fail to respond to immunotherapies or relapse following treatment^1^. Recent efforts have focused on using CRISPR knockout or activation screening to identify targets that enhance T cell effector function and augment immune killing capability^2^. Nevertheless, overcoming resistance by manipulating a single gene remains challenging due to the compensatory effects of other genes. Combination therapies have been proposed as a promising strategy to overcome tumor resistance to monotherapy when single agents are ineffective^3^.

Paralogs, genes that originate from the same ancestors and share similar functions, often work in concert to promote cell survival. Numerous focused studies have utilized genomic data and pooled screening techniques to explore how targeting paralogous genes can impact cancer cell viability^4–6^. Recent research has shown that targeting these genes simultaneously in cancer cells can enhance the effectiveness of immunotherapy by preventing the cells from evading T-cell-mediated destruction. For instance, Park *et al*. demonstrated that treating mice with both anti-PD-L1 and anti-PD-L2 antibodies significantly reduced tumor growth in specific mouse models^7^. Additionally, signals from both *PARP-1* and *PARP-2* are essential for initiating an effective T-cell response against breast cancer in mice^8^. While pooled CRISPR screening is popular for identifying single genes that modulate specific responses, including those relevant to immunotherapy, few efforts have focused on double-knockout screens. Double-knockout screens aim to identify novel combinations of genes that lead to synthetic lethality and boost T-cell function^5,9,10^. However, a comprehensive, high-throughput strategy to screen all potential paralog pairs related to immunotherapy is challenging because a very large screening library, that includes hundreds of thousands of gene combinations, would be required. To solve this problem we have developed a computational approach to identify the most promising paralog gene combinations, which can then be used to build a focused and manageable library for experimental validation.

Here we used an sgRNA sets enrichment method to identify cancer-intrinsic paralog pairs that enhance T cell killing from prior pooled screening data. We show that our method identifies novel synergistic paralog pairs that are successfully validated experimentally. We have also constructed an ensemble learning XGBoost classifier to predict true-positive paralog pairs enhancing immunotherapy by incorporating features such as gene characteristics, sequence and structural similarity, protein–protein interaction networks, and gene coevolution data. To enhance reliability of the XGBoost classifier predictions, we conducted further experimental validations on the highest-ranked paralog pairs identified. This study is expected to uncover novel therapeutic targets and inspire new combination immunotherapies to aid cancer treatment.

## Results

### Identifying cancer-intrinsic paired paralogs that synergistically enhance T cell-mediated killing using CRISPR screen data

A large proportion of genes have paralogs that perform redundant functions. Inactivation of functionally important paralog pairs can lead to synthetic lethality as well as more efficient modulation of biological pathways^9,11^. Unfortunately the identification of these paralog pairs is limited in single-gene perturbation screens due to compensation between paralogous genes^12^ and dual gene screening is often unfeasible due to the large size of CRISPR libraries required to encompass all potential combinations of paralogous gene pairs (**Fig. 1A**). To overcome this limitation we proposed an unbiased computational approach to filter and predict functional gene pairs from prior published genome-wide CRISPR screen data.

**Figure 1.**
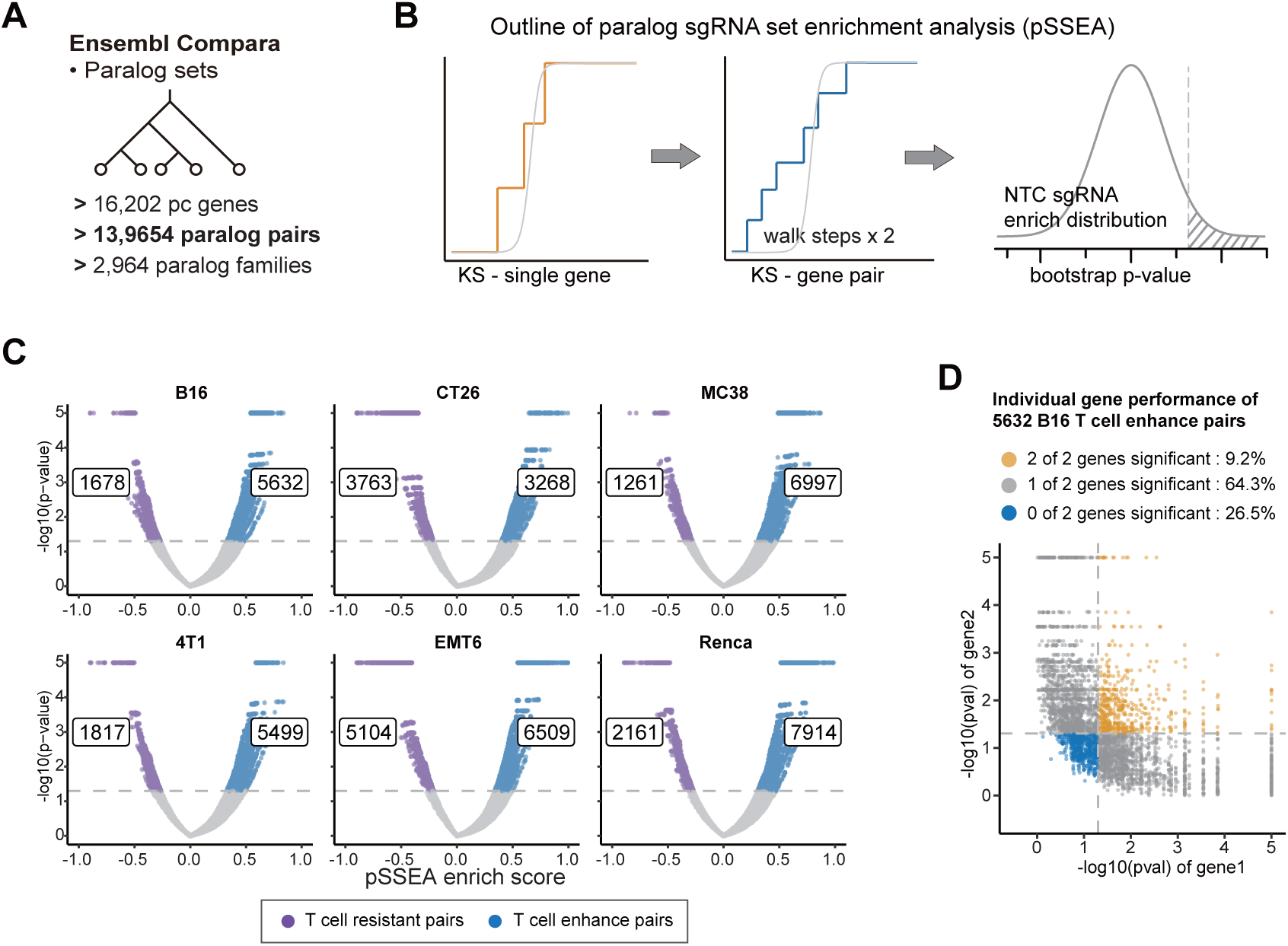
Identification of paralog pairs that enhance T cell killing using pSSEA. (A) Protein-coding paralogous genes retrieved from Ensembl Compara database using bioMart (B) Workflow for identifying paralog pair hits using the paralog sgRNA Set Enrichment Analysis (pSSEA). (C) Identification of paralog pairs that enhance T cell killing in six cancer models using pSSEA. CRISPR screen data was from Lawson et al. 2017. The volcano plot of pSSEA outputs, with the x-axis representing the enrichment score and the y-axis showing the negative logarithm (base 10) of the p-value from pSSEA, is shown. (D) Scatter plot illustrating the characteristics of identified T cell enhancing paralog pairs using a single gene enrichment model. The x-axis and y-axis represent the negative logarithm (base 10) of the p-values from the enrichment results of gene1 and gene2, respectively.

We initially attempted to search for immunotherapy-aiding paralog pairs using conventional MAGeCK results in Lawson’s CRISPR screen dataset^13^, which was originally aimed at identifying genes involved in evading T cell-mediated cytotoxicity across six murine tumor cell lines. Based on a strict dual-hit genes criterion, we observed few or no suitable pairs identified (**Fig. S1A**); however, relaxing the criteria to include single hit genes leads to a significant increase in uncertain pairs. To overcome this we developed an enrichment-based methodology named Paralog sgRNA Set Enrichment Analysis (pSSEA), which integrates the sgRNAs of gene pairs to enhance the identification of high-potential paralog pairs. We provide a schematic of the pSSEA algorithm in **Fig. 1B**. Briefly, the process begins with the results of differential expression analysis (using DESeq2, MAGeCK, etc.) as input. The pSSEA evaluates the enrichment score employing a Kolmogorov-Smirnov (KS)-like random walk statistic, a method similar to those used in GSVA^14^ and ASSESS^15^. The detailed steps include:

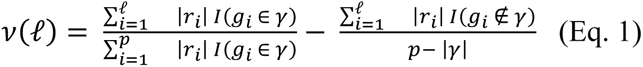

where *r_i_* is the rank-ordered sgRNAs, *γ* is the sgRNAs set that belongs to the paralog gene pair, *I*(*g_i_* ∈ *γ*) is the indicator function on whether the i-th sgRNA is in paralog sgRNAs set *γ*, |*γ*| is the number of sgRNAs belonging to the gene pairs, and p is the number of total sgRNAs in the library.

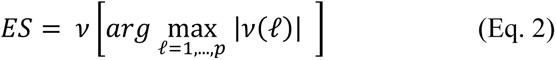

The enrichment scores for paralog pairs were calculated by identifying the maximum deviation from zero during a random walk analysis.

To assess the significance of enrichment, we created an empirical null distribution by randomly selecting an equivalent number of non-targeting control (NTC) sgRNAs and repeating this process 10,000 times to ensure robustness. The significance of enrichment for each gene pair was then determined by comparing their sgRNA distributions against both the positive and negative tails of the NTC null distribution.

We re-analyzed Lawson’s CRISPR screen dataset^13^ to identify cancer intrinsic paralog pairs that enhance T cell-mediated killing function using the pSSEA method. From the EnsemblCompara database (Ensembl, v102), we obtained a total of 139,654 protein-coding gene paralog pairs, covering 16,202 protein-coding genes, as shown in **Fig. 1A**. First, DESeq2 was used to quantify the selective depletion of sgRNAs within tumor populations under immune selection from co-cultured OT-1 T cells. Subsequently, using the pSSEA framework, we identified paralogous gene pairs that potentially influence T cell-mediated cytotoxicity, based on the differential sgRNA results obtained from DESeq2. Our analysis identified 7,310 significant paralog pairs (5.2% of 139,654) with p-values below 0.05 in B16 melanoma cell data. Among these, 5,632 pairs were positively enriched suggesting vulnerability to T-cell killing, and 1,678 pairs were negatively enriched suggesting resistance to T-cell killing (**Fig. 1D**). Using these same methods we identified paralog pairs associated with vulnerability and resistance to T cell killing for CT26, MC38, 4T1, EMT6 and Renca cell lines, as shown in **Fig. 1C**.

To investigate the properties of paralog pairs identified as enriched through pSSEA analysis, we assessed individual gene enrichment using pSSEA in single-gene mode. Among the 5,632 pairs associated with vulnerability to T-cell killing in B16 melanoma cell, 3,620 pairs (64.3%) were single-hit, with only one gene showing significance; 520 pairs (9.2%) were dual-hits, with both genes significant; and the remaining 1,492 pairs (26.5%) showed no hits of either of the genes within the identified paralog pair (**Fig. 1D** and **Table S1)**. Notably, we observed that over a quarter of the paralog pairs exhibited a synergistic effect despite no hits at the single-gene level, indicating a broad spectrum of potential targets that might be overlooked by obtaining gene pairs according to individual gene results (**Fig. S1A).**

### Validation of synergistic cancer-intrinsic paralog pairs that together boost T cell killing

For the experimental validation of the synergistic effect of gene pairs in which neither individual gene reached significance, we selected the most promising paralog pairs using the following criteria: 1) the genes within the paralog pair must show significance in combination only (neither gene is individually significant) in B16 melanoma cells, due to the availability of murine cell lines, and 2) the paralog pairs must show significant enhancement of T cell killing, as assessed by pSSEA, in at least three out of the five screened cancer cell types, excluding B16 (**Fig. 2A** and **Table S2**). Ultimately, four paralog pairs, *Rbm45+Rbms2*, *Ppp2r2a+Ppp2r2d*, *Elf2+Etv6*, and *Adam10+Adam15* were selected for further experimental assessment of their potential to enhance T cell killing efficiency (**Fig. S1B-F**).

**Figure 2.**
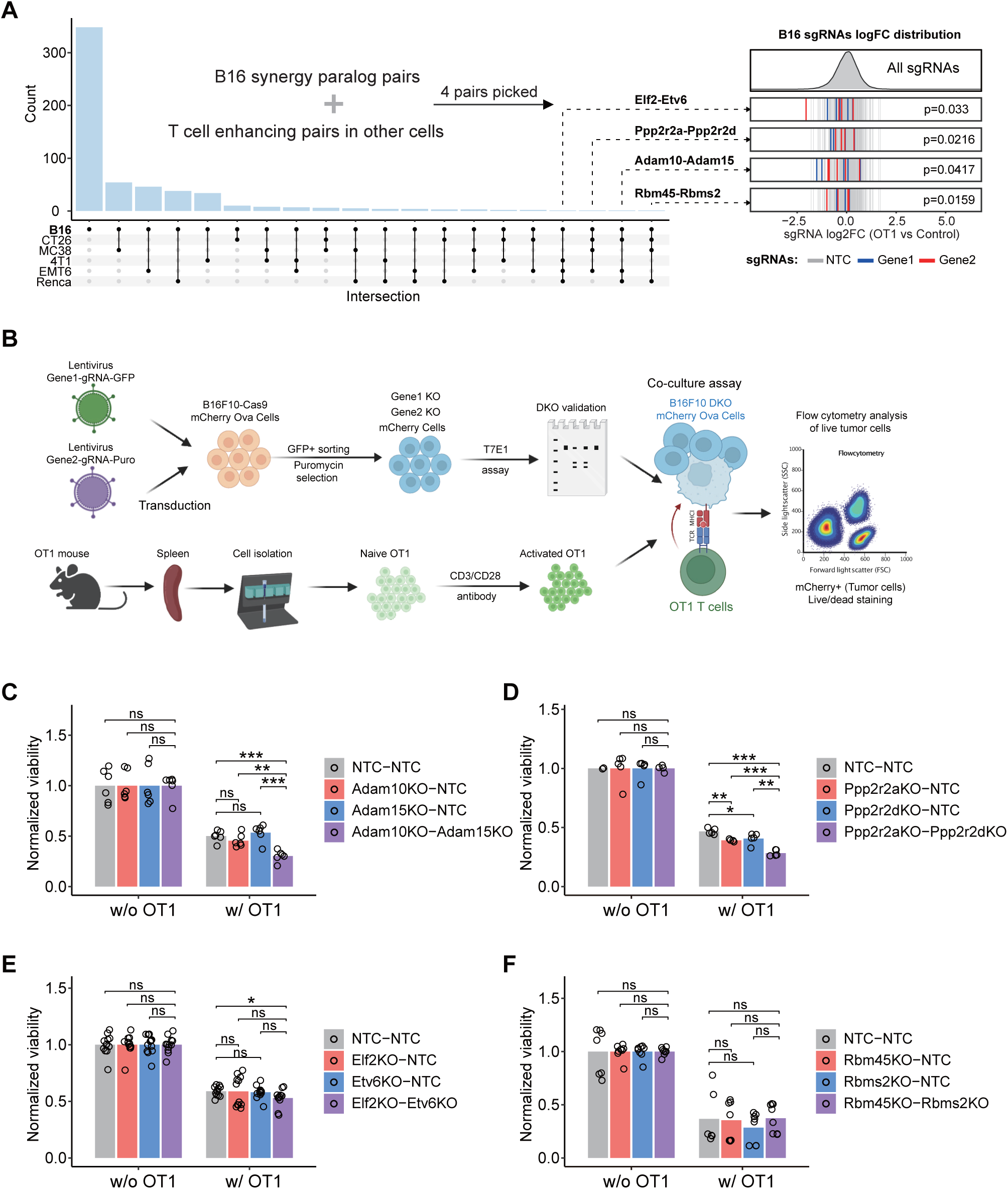
Synergistic paralog pairs that together boost T cell killing function in B16F10 tumor model. (A) The T cell synergistic paralog pairs identified in the B16F10 melanoma model and the intersection across the other 5 tumor models. T cell enhanced paralog pairs were identified using pSSEA, and synergistic pairs were denoted as pairs where neither of the two genes could reach significance using a single gene enrich approach. Four paralog pairs were identified in 4 out of 6 tumor models. Left panel: upset plot of overlap synergistic paralog pairs identified from B16F10 melanoma model that were also significant in the other 5 cancer cell models in Lawson study. (B) Experimental design for the validation of selected paralog pairs predicted to enhance T cell killing. Naive CD8a+ T cells were isolated from the spleen of OT-I mice and stimulated with anti-mouse CD3/CD28 antibody for 2 days. Then B16F10-Cas9-mCherry-Ova gene knockout cells were co-cultured with OT-I T cells at a E:T=1:1 ratio. Tumor cell killing were tested at 24 hours by flow cytometry. (C) Bar chart of mean viability of B16F10 cells with *Adam10* and *Adam15* single or double gene knockout/s following culture with or without OT-1 T cells. Individual data points are shown as black dots. The p values come from two-sided heteroscedastic t tests (ns, p>0.05; *, P ≤ 0.05; **, P ≤ 0.01. ***, P ≤ 0.001). (D) Bar chart of mean viability of *Ppp2r2a* and *Ppp2r2d* knockout/s B16F10 cells following culture with or without OT-1 T cells. (E) Bar chart of mean viability of *Elf2* and *Etv6* knockout/s B16F10 cells following culture with or without OT-1 T cells (F) Bar chart of mean viability of *Rbm45* and *Rbms2* knockout/s B16F10 cells following culture with or without OT-1 T cells.

To perform the paired gene knockout, we utilized the same gRNAs from the original gRNA-library pool used in the CRISPR screen. For the first gene within the paralog pair, the targeting gRNA was cloned into a GFP-expressing construct, while for the second gene within the paralog pair, the targeting gRNA was inserted into a puromycin-expressing construct (**Fig. 2B**). To evaluate the synergistic function of each gene pair, we established four experimental groups with targeted knockouts : non-targeting control (NTC) paired with NTC (NTC-NTC), gene1 knockout (KO) paired with NTC (gene1KO-NTC), gene2 KO paired with NTC (gene2KO-NTC), and a double knockout (DKO) of both genes (gene1KO-gene2KO). To generate cells with these specific gene targets, we performed dual transduction of B16F10-Cas9-mCherry-OVA tumor cells, which express the OVA antigen, using lentiviruses carrying the gRNAs with GFP or puromycin markers. GFP-positive cells were sorted and then selected for puromycin-resistant cells to obtain either single or double knockout cells. Subsequently, gene editing was verified and knockout efficiency was evaluated using the T7E1 assay. To investigate the role of the paired genes in T cell-mediated cytotoxicity, we co-cultured B16F10-Cas9-mCherry-OVA tumor cells from the four experimental groups described above with OT-I T cells, whose TCR recognizes the SIINFEKL peptide presented by Ova antigen-expressing tumor cells, and assessed tumor cell survival.

In these co-culture experiments we found that simultaneous inactivation of *Adam10* and *Adam15* significantly increased T-cell-mediated tumor cell death, while single knockouts of *Adam15* or *Adam10* didn’t affect T-cell activity, (**Fig. 2C** and **Fig. S1G**). Similarly, *Ppp2r2a+Ppp2r2d* double knockouts displayed significantly greater vulnerability to T-cell cytotoxicity, whereas individual knockouts of *Ppp2r2a* or *Ppp2r2d* showed only modestly enhanced T-cell killing (**Fig. 2D** and **Fig. S1H**). The *Elf2* and *Etv6* combination showed significantly enhanced T cell kill compared to the NTC control, but not compared to the single gene *Elf2* and *Etv6* knockout cells (**Fig. 2E and Fig. S1I**). The *Rbm45/Rbms2* pair showed no discernible impact on T-cell efficiency in both single and double knockout cells compared to the NTC group (**Fig. 2F** and **Fig. S1J**). In total, three out of four predicted paralog pairs were successfully confirmed to exhibit increased T cell vulnerability, highlighting previously overlooked synergistic gene pairs. These findings also provide fundamental experimental evidence for the accuracy of using pSSEA method to effectively identify paralog pairs that enhance T-cell killing efficacy.

### Building an ensemble XGBoost classifier to predict paralog pairs that enhance T cell-mediated cytotoxic killing

Using a CRISPR screen dataset we have demonstrated that paralog pairs could act synergistically to enhance T cell mediated cytotoxicity. However, this CRISPR screen dataset was derived from *in-vitro* co-culture of cancer cell lines with cytotoxic T cells, an assay with significant limitations and potential experimental biases. We therefore developed a system for deeper characterization of paralog pairs and prediction of their potential for enhancing cancer immunotherapy. Utilizing T cell enhancing paralog pairs identified in B16F10 melanoma data with pSSEA as true positives and those insignificant as false positives, we compiled 32 features representing various aspects of paralogous gene interactions. These features include sequence characterization, expression abundance and correlation, shared protein-protein interactions (PPI), complex membership, co-evolution, tumor microenvironment (TME) associations, perturbation similarities, and combined survival associations in cancer patients (**Fig. 3A** and **Table S3**). Then, an ensemble classifier was trained, utilizing the identified 32 features, to predict potential paralog pairs. To ensure the quality of model training and testing, the paralog pairs were filtered using several criteria: paralog pairs identified as T cell enhancing pairs or not significant pairs in B16 data; sequence BLAST identity larger than 30% with an e-value < 1e-5; with measurements in structural similarity, synthetic lethality and survival associations; and paralog pairs with family sizes of 20 or fewer to prevent overrepresentation of specific ortholog families. A total of 3,204 paralog pairs were selected from the initial set of 139,654 paralog pairs. Furthermore, 80% of these 3,204 pairs were randomly assigned to the training dataset (n = 2,547), which includes negative pairs (n = 2,274) and positive pairs (n= 273). The remaining 20% were saved for internal testing later (**Fig. S2A-B)**. Prior to building the ensemble model, our analysis revealed that certain individual features exhibited a stronger predictive effect in the training data. The top influential feature was gene expression abundance with the highest ROC AUC (area under receiver operating characteristic curve) values, indicating that gene expression itself could be a critical indicator of a paralog pair(**Fig. 3B**). Notably, the features ranked second to fifth were four protein-protein-interaction (PPI) -network features (**Fig. 3A**), highlighting the crucial role of gene networks in determining whether two genes could interact in an immunotherapy context. Synthetic lethality was identified as a mid-level feature, suggesting that the collaborative roles of paralogs of cancer cells in response to immune context might differ from their roles in cell viability.

**Figure 3.**
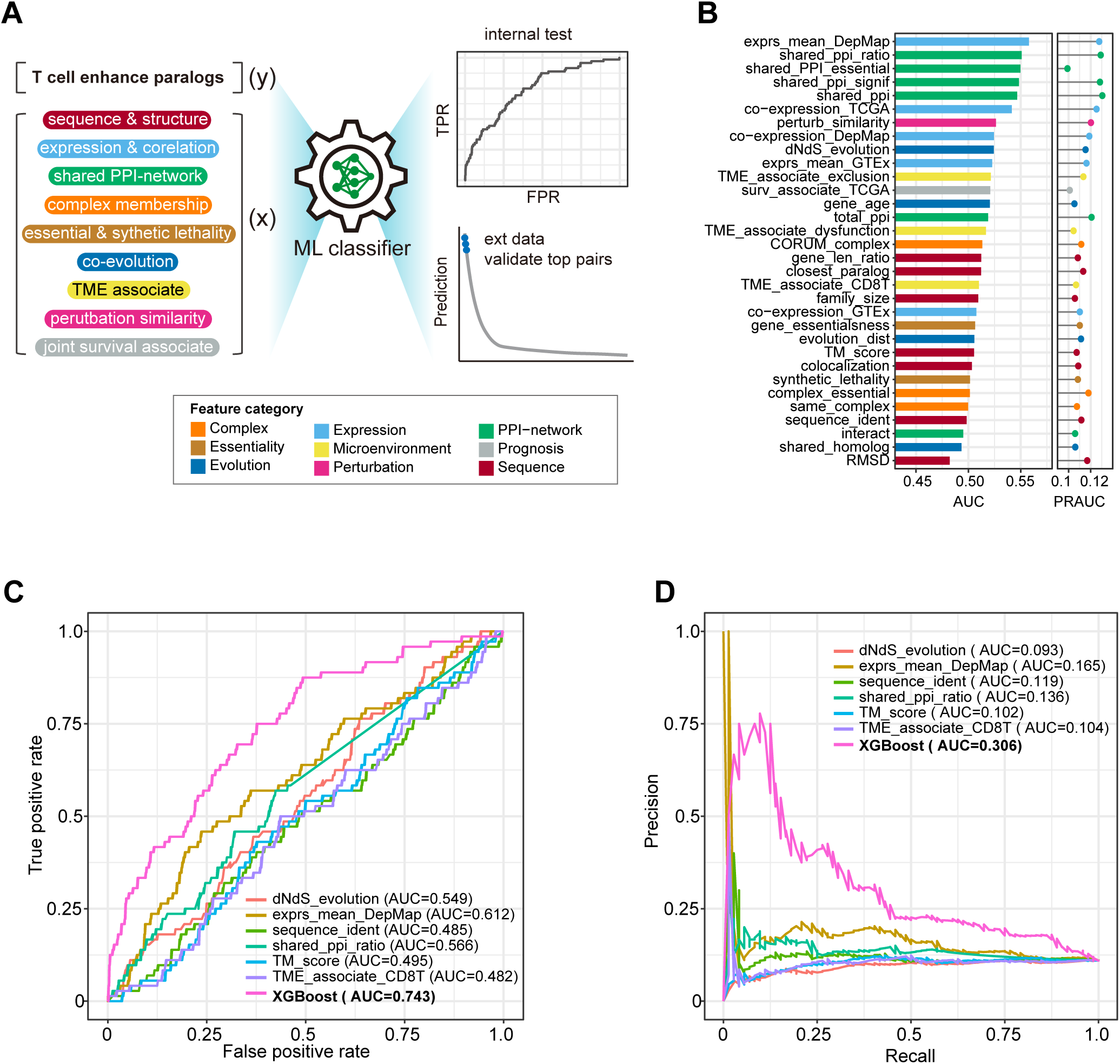
Development and evaluation of XGBoost classifier for predicting T cell enhancing synergistic paralog pairs. (A) Schematic for constructing a machine learning classifier to predict T cell enhancing paralog pairs. (B) AUROC and AUPRC measures of 32 paralog features for predicting paralog pairs that boost T-cell killing. (C) ROC curves of XGBoost and top individual features in the internal test dataset. The XGBoost classifier outperformed the traditional individual features. (D) Precision recall curves of the XGBoost classifier and individual features.

Next, XGBoost classifier, an ensemble boosted tree learner, was trained, utilizing 5-fold cross-validation for hyperparameter tuning. We then applied the optimal parameters to re-train the classifier using the entire training dataset (details in Methods). Variable importance analysis revealed that complex, colocalization and PPI features were the top-ranking features (**Fig. S2C**). We then evaluated the XGBoost classifier’s performance using testing data. Notably, the XGBoost classifier significantly outperformed all single features used as a baseline for comparison (**Fig. 3C**). Specifically, the classifier achieved an AUC of 0.743, surpassing the best-performing individual feature, average paralog gene expression in cancer cell lines, which had an AUC of 0.612. Considering the imbalanced nature of the training dataset, with more negative pairs than positive ones, we also assessed the classifier’s performance using the Precision-Recall Curve (PRC; **Fig. 3D**). Similarly, XGBoost classifier showed better performance compared to all the individual features. Overall, the results underscore the enhanced predictive capability of our XGBoost classifiers in effectively capturing interactions between features and non-linear relationships, thereby enhancing T cell killing potential.

Furthermore, we evaluated the classifier’s performance on paralog pairs in cells other than B16F10 cells. ROC analysis revealed that the classifier trained on B16F10 cell data could not predict paralog pairs enhancing T cell killing in other cell types, except for CT26 colon cells (**Fig. S3A-E**), indicating inherent heterogeneity among cancer cell models.

### Interpreting the prediction of immunotherapy favorable paralog pairs from the machine learning classifier

Using the XGBoost classifier, we comprehensively analyzed the probability of enhancing T cell cytotoxicity for additional paralog pairs outside the training and internal testing sets. By filtering with criteria including measurements in structural similarity, synthetic lethality, survival associations, and excluding pairs from the training and testing sets, we obtained 15,414 paralog pairs out of the total 139,654 pairs for prediction.. Each paralog pair was calculated for its potential impact, assigned a probability score, and ranked accordingly. The SHapley Additive exPlanations (SHAP) tree explainer method assessed the contribution of each feature to the final predictions.

The paralog pairs were ranked by their prediction scores, and subsequent pathway enrichment analysis revealed that genes in the top 5% of paralog pairs were significantly enriched in functions related to protein phosphorylation and ubiquitination modification (**Fig. S4A**). The top three ranked predicted paralog pairs were: *Rapgef1+Papgef2*, *Cct8+Cct3*, and *Syk+Itk* (**Fig. 4A** and **Table S3**). Notably the expression of *Syk* in cancer cells has shown to be associated with both tumor promotion and suppression^16^. *Itk*, has been reported to enhance immune checkpoint blockade response in solid tumors^17^ and *Rapgef1* was identified as essential for melanocyte growth through a CRISPR proliferation screen^18^. Analysis of the Cancer Genome Atlas (TCGA) cohort revealed that four genes (*Rapgef1*, *Rapgef2*, *Itk* and *Syk*) had mutation rates exceeding 1% in pan-cancer cohort. At the individual cancer type level, four genes (*Cct3*, *Rapgef1*, *Rapgef2* and *Syk*) had mutation rates exceeding 5% in high mutated endometrial carcinoma, while in melanoma three genes (*Rapgef1*, *Rapgef2*, and *Syk*) had mutation rates exceeding 3%, with *Itk* having a rate of 7.3% **(Fig. 4B**). The SHAP profile revealed differential feature attributions among the top pairs. For example, sequence identity and paralog family size showed negative contributions to the prediction in *Rapgef1+Rapgef2* (**Fig. 4C**) and *Itk+Syk* (**Fig. 4D**), but positive contributions in Cct3+Cct8 (**Fig. S4B**).

**Figure 4.**
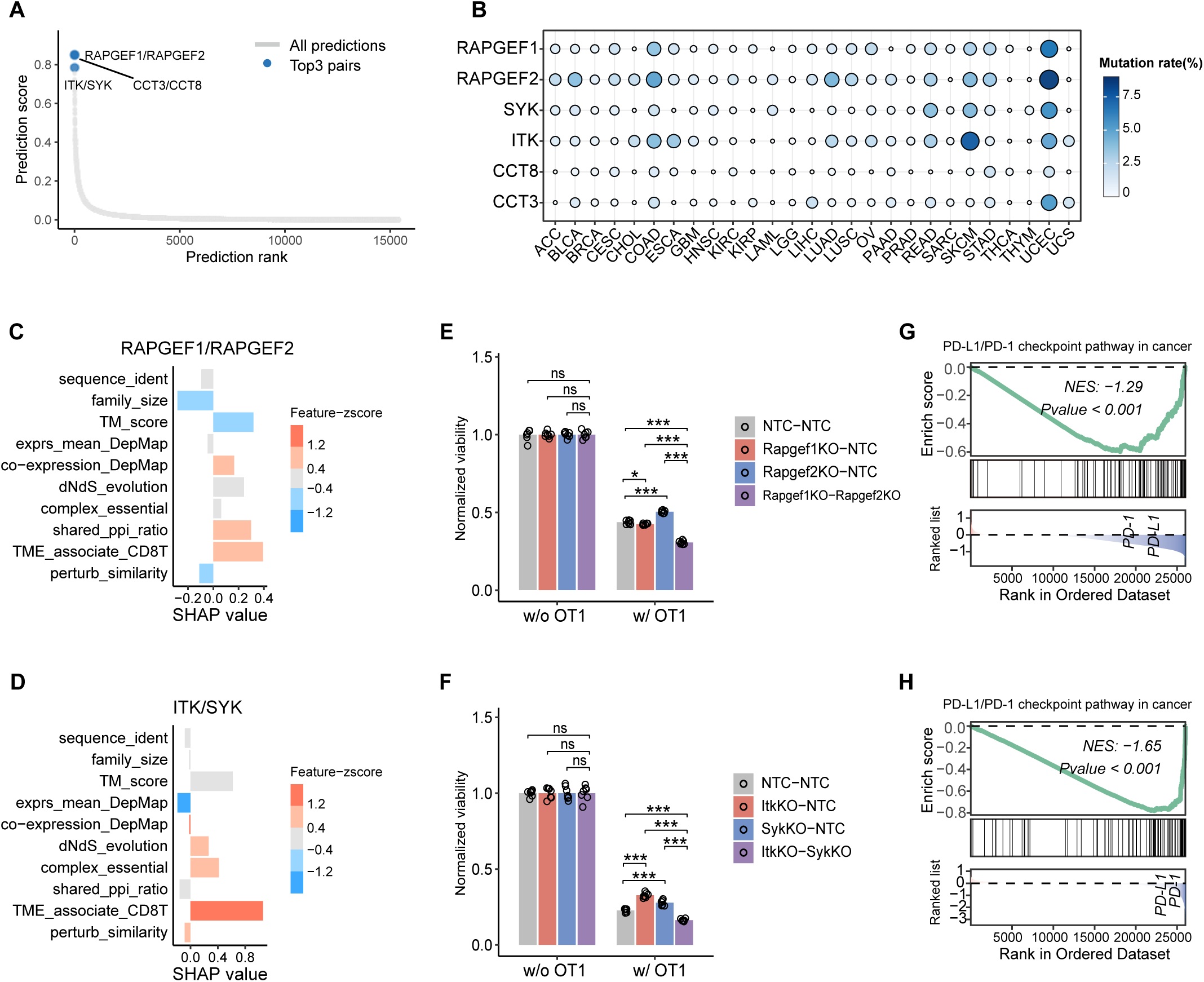
Visualization and validation of top 3 predicted paralogs from XGBoost classifier. (A) Paralog pairs with the top 3 prediction scores are displayed in a scatter plot. (B) Mutation profile of top 3 paralog pairs *RAPGEF1/RAPGEF2*, *SYK/ITK* and *CCT8/CCT3* in the Cancer Genome Atlas (TCGA) cohorts. (C, D) SHAP profile for *RAPGEF1/RAPGEF2* (C) and *SYK/ITK* (D). (E, F) Bar chart of mean viability of *Rapgef1/ Rapgef2* (E) and *Itk/Syk* (F) knock-out treated B16F10 melanoma cells following culture with or without OT-1 T cells. The p values come from two-sided heteroscedastic t tests. (G, H) Gene sets enrichment analysis (GSEA) revealed that patients with low expression of *RAPGEF1/RAPGEF2* (G) and *SYK/ITK* (H) were associated with downregulation of PD-L1/PD-1 signaling pathway. The GSEA was performed using the differential expression output between patients with low (<25%) expression of both paralog genes versus all remaining patients.

To evaluate the synergistic effect of the paralog genes identified by XGBoost classifier, we conducted DKO-B16F10-OT1 T cell co-culture assays. Knocking out *Rapgef1* alone did not affect T cell-mediated tumor cell killing, while knocking out *Rapgef2* alone increased tumor cell resistance to T cell killing. However, a combined knockout of *Rapgef1* and *Rapgef2* significantly improved T-cell efficiency in killing tumor cells (**Fig. 4E** and **Fig. S4C**). A similar pattern was observed with the *Syk+Itk* gene pair, knocking out individual genes increased resistance to T cell-mediated killing, but a double knockout enhanced T cell killing activity (**Fig. 4F** and **Fig. S4D**). In contrast, double knockout of the *Cct8+Cct3* gene pair impaired T cell killing ability against tumor cells (**Fig. S4E** and **S4F**).

To further investigate why *Cct3/Cct8* might fail while the other two pairs work, we performed gene set enrichment analysis (GSEA) on the TCGA-SKCM data, comparing patients with low expression in both paired genes (<=25%) to those with high expression (>=75%). We observed significant downregulation of the PD-L1/PD-1 signaling pathway in melanoma patients with low *Rapgef1+Rapgef2* (**Fig. 4G**) and *Itk+Syk* (**Fig. 4H**) expression. Downregulation of the PD-L1/PD-1 signaling pathway augments T-cell killing by interrupting inhibitory signals that suppress T-cell activity. Conversely, patients with low *Cct3+Cct8* expression (**Fig. S4G**) showed insignificant upregulation of this pathway compared to others. This finding suggests that the failure of *Cct3+Cct8* to enhance T cell cytotoxicity may be due activation of other immunosuppressive pathways and the elevated expression of inhibitory molecules.

## Discussion

Although numerous CRISPR screen studies aim to identify novel cancer cell targets for enhancing cancer immunotherapies, few have focused on functional gene pairs, thus limiting the development of combination strategies. To address this, we developed a computational enrichment-based approach, pSSEA, to identify potential paralog gene pairs using genome wide CRISPR screen datasets by combining sgRNAs from two paralogs. We demonstrated that pSSEA enables the identification of cancer-intrinsic paralog pairs that can synergistically enhance T cell killing. Subsequently, we constructed an ensemble-learning XGBoost classifier to predict additional cancer-intrinsic-immunotherapy-aiding paralog pairs and experimentally tested the top predictions. Notably, all analyses in this study were based on CRISPR screen data. We observed that only a small subset of the screen-derived T cell-enhancing paralog pairs demonstrated a combined effect in predicting ICB responses when using patient-derived transcriptomic data (data not shown). This suggests that these two data types may capture different aspects of immunotherapy-related signals.

In summary, we envision our work providing novel methods to leverage genome-wide screen data for selecting combination targets in cancer immunotherapy. We believe the computational framework in our study could be adapted to prioritize non-paralogous protein-coding gene pairs, broadening its applicability beyond paralogs.

## Acknowledgments

We thank all members in Chen laboratory, as well as various colleagues in Yale Genetics, SBI, CSBC, MCGD, Immunobiology, BBS, YCC, YSCC, and CBDS for assistance and/or discussions. We thank various Yale Core Facilities for technical support.

S.C. is supported by Cancer Research Institute Lloyd J. Old STAR Award (CRI4964), NIH/NCI (R33CA281702), DoD (W81XWH-21-1-0514, HT9425-23-1-0472, HT9425-23-1-0860), Alliance for Cancer Gene Therapy (ACGT), and Pershing Square Sohn Cancer Research Alliance. CD is supported by Boehringer Ingelheim Biomedical Data Science Fellowship. NV is supported by American Board of Radiology’s B. Leonard Holman Research Pathway Fellowship and ASTRO Seed Grant.

## Method

### Paralog pairs information

Information on protein-coding paralog pairs was obtained from Ensembl Compara database via biomaRt (hsapiens_gene_ensembl, version 102). Duplicated paralog pairs were filtered out. A total of 139,654 protein-coding gene paralog pairs, covering 16,202 protein-coding genes, were collected for the analyses in this study.

### Collect features for paralog pairs. Sequence similarity

Amino acid sequence of paralog genes were obtained from Ensembl (Release 102)^19^. Paralog sequence similarities were measured using BLASTP command line tools^20^. The longest translated peptide was used for genes with multiple transcripts. Gene pair sequence identity, paralog family size, and whether they are the closest pair in the same gene family were included as sequence-related features for paralog pairs.

We also used high-resolution structures PDB files from https://alphafold.ebi.ac.uk/^21^. The structural similarity of paralogous proteins was measured using TM-score software^22^. The template modeling score (TM-score), and root mean square deviation (RMSD) were included as additional sequence-related features.

### Shared gene network, complex, and co-localization

We collected shared protein-protein interaction (PPI) features to characterize the role of paralogs in the gene interaction network, including total PPIs, shared PPIs, and two statistical measures: the Jaccard index and Fisher exact test -log10(p-value) for shared PPIs. Gene subcellular location data were obtained from the Human Protein Atlas (version 22.0, file: subcellular_location.tsv). The co-localization of paralogous genes was also measured using the Jaccard index. The complex membership of paralog pairs and gene-protein complex membership data were obtained from CORUM^23^ via https://maayanlab.cloud/Harmonizome/.

### Gene co-evolution

Gene ages were obtained from ProteinHistorian (https://proteinhistorian.docpollard.org/)^24^, and we used the average age of the two paralogs as the age for the pair (wagner age reconstruction algorithm). The conservation level of an individual gene was calculated as the number of species with orthologs (BLASTP e-value < 1-e-5, identity >30%) out of 269 species from Ensembl genomes (version 102). The shared homologs of a gene pair was calculated as the number of species having both gene homologs. The ratio of the number of non-synonymous substitutions per non-synonymous site to the number of synonymous substitutions per synonymous site (Ka/Ks) for gene pairs was calculated using the codeml program in the PAML package (version 4.10.6)^25,26^. We adopted the phylogenetic distance method developed by Tabach *et al*. to measure the co-evolution of a pair of genes^27^. A non-negative matrix factorization (NMF)-derived distance considering phylogenetic tree information was calculated using the phylogenetic profiles of the paralogous genes^28,29^.

### Essentiality and synthetic lethality

Paralogs with available evidence of essentiality from S. pombe and S. cerevisiae ortholog data from OGEE v3 were marked as essential pairs^30^. We adopted Kegel’s method to identify synthetic lethality pairs^6^ using DepMap CRISPR data (https://depmap.org/portal<u>, 23Q2 release</u>). Paralog pairs were considered synthetic if the A1 gene was essential gene (Chronos score^31^ < −0.6 in 1% or 10% of cell lines), and its dependency was significantly associate with A2 loss status with p-value less than 0.05.

### Co-expression pattern

Gene pair co-expression was measured using Spearman correlation coefficient ρ at multiple levels: cancer cell lines, normal tissues, and cancer scenarios. Cancer Cell Line Encyclopedia (CCLE) gene expression data were downloaded from DepMap^32^. Tissue-specific gene expression profiles from healthy donors were downloaded from Genotype-Tissue Expression (GTEx, https://gtexportal.org/) public releases^33^. Cancer-specific gene expression profiles for 33 cancer types were downloaded from the TCGA GDC portal (https://portal.gdc.cancer.gov/)^34^.

### Perturbation similarity

Due to the limited number of gene perturbation experiments in Connectivity Map (L1000, https://clue.io/data/)^35^, we conducted pseudo perturbation using TCGA mRNA sequence data to mimic the transcriptomic gene expression changes associated with single gene manipulation. Practically, the differential expression profile of a specific gene inhibition was calculated as the log fold change in gene expression between patients with expression levels in the lowest 25^th^percentile versus those in the highest 75th percentile of the query gene. The perturbation similarity of paralogous genes was measured using recovery AUC with the paralog gene differential expression profiles.

### Microenvironment associations

The tumor microenvironment plays an important role in cancer progression and treatment outcome. We attempted to uncover features quantifying the association of the paralogs with multiple microenvironment indicators. CD8+ T cell infiltration was measured by the ssGSEA method^14^. We calculated T cell exclusion and dysfunction scores using another software, TIDE algorithm developed by Jiang *et al*^36^. The association with gene pairs and microenvironmental characteristics was calculated by Kendall’s rank correlation test^37^.

### Survival associations

To determine gene pairs with a synergistic effect on survival outcomes, we calculated the survival associations to assess whether a paralog pair acts in concert to influence cancer prognosis. First, patients were categorized into three groups based on the expression levels of the paired paralog genes: groups were designated as 1 for both genes showing low expression, 3 for both showing high expression, and 2 for any other expression combination. The survival association for paralog pairs was then calculated using the formula:

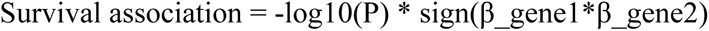

In this formula, P refers to the p-value from the Cox regression analysis of the paralog group, adjusted for sex and age, while β_gene1 and β_gene2 represent the exponentiated coefficients of the individual genes from the Cox proportional hazards model. Clinical data and genomics data for TCGA-SKCM were downloaded from GDC portal (https://portal.gdc.cancer.gov/)^38^.

### XGBoost ensemble classifier

The popular supervised-learning algorithm, XGBoost – a gradient boosting ensemble learner with regularization parameters was used in this study (implementation in the XGBoost package)^39^. We performed a grid search based on 5-fold cross-validation of our paralog pair dataset to explore the space of potential hyper-parameters for the XGBoost classifier. The parameters used in the final model are as follows: objective = reg: logistic, n_estimators = 400, max_depth = 10, min_child_weight = 1, min_split_loss = 0.1, subsample = 1, colsample_bytree = 1, learning_rate = 0.01, and random_state=8 (for reproducibility). All parameters not listed used default settings.

### Performance evaluation

The ROC AUC and PRC AUC for the feature values as well as the classifiers were computed to evaluate the performance of individual features and trained models.

### Lentivirus purification

For lentivirus production, 20 μg of lenti-U6-sgRNA-EFS-GFP-WPRE, or 20 μg of lenti-U6-sgRNA-EFS-Puro-WPRE, 10 μg of pMD2.G, and 15 μg of psPAX2 were co-transfected into LentiX-293 cells plated in a 150 mm-dish at 80-90% confluency using 130 μg polyethyleneimine (PEI). 6 hours later, the media was replaced with fresh DMEM+10%FBS. Virus supernatant was collected 48 h post-transfection and centrifuged at 1,500 g for 15 min to remove the cell debris. The virus supernatant was concentrated by Lenti-X concentrator at 4°C for 30 minutes (1 volume of Lenti-X concentrator with 3 volumes of supernatant), followed by centrifugation at 1,500 x g for 45 minutes at 4°C. Finally, the virus was resuspended in DMEM, aliquoted, and stored at −80°C.

### Generation of single-gene-knockout or double-gene-knockout cells

B16F10 cells stably expressing Cas9 were generated by transducing B16F10 cells with lentiviral EF1a-NLS-Cas9-2A-Blast-WPRE, followed by 5 days of selection under 20 μg/ml blasticidin. B16F10-Cas9 cells were further transfected with lentiviral EF1a-mCherry-2A-OVA-WPRE and sorted for mCherry positive cells to generate B16f10-Cas9-mCherry-OVA cells. The two gRNAs targeting each gene within the paralog pair were transduced into the B16F10-Cas9-mCherry-Ova cells simultaneously. Guide RNA virus infected cells were cultured at 37°C for more than 24 h, and then GFP positive cells were sorted and selected under 5 μg/ml puromycin 3 days to generate the double knockout cells.

### T7E1 assay

T7E1 was used to estimate gRNA cutting efficiency. In brief, gDNA was extracted by using Genomic DNA prep with Quick Extract (QE) buffer. PCR amplification of the genomic regions flanking the crRNAs was performed using the primers listed in supplementary **Table S2** Using Phusion Flash High Fidelity Master Mix (Thermo Fisher Scientific), the thermocycling parameters for PCR were 98 °C for 1 minutes, 35 cycles of (98 °C for 1 second, 60 °C for 5 seconds, 72 °C for 15 seconds) and 72 °C for 2 minutes. The PCR amplicons were then used for T7E1 assays according to the manufacturer’s protocol.

### Naïve OT-I CD8a + T cell isolation and co-culture assay

The Naive OT-I CD8a+ T Cells were isolated using the Naive CD8a+ T Cells isolation Kit (Miltenyi Biotec) following the manufacturer’s instructions. Naive CD8a+ T cells were isolated from the spleen of OT-I mice and stimulated with anti-mouse CD3/CD28 antibody for 2 days. For the tumor cell and OT-I T cell co-culture assay, B16F10-Cas9-mCherry-Ova gene knockout cells were co-cultured with OT1 cells at E:T=1:1 ratios. Tumor cell killing were tested at 24h hours by flow cytometry. The LIVE/DEAD Near-IR was diluted at 1:1000 to identify the dead cells. Tumor cells were identified as mCherry positive cells.

### Data availability

The interactively web application of this study is deployed at https://sidichenlab.shinyapps.io/imparalog for free visiting and querying. All the processed input data, analysis output data can be downloaded from the website.

### Code availability

The source code for processed data and modeling is available under GitHub at https://github.com/cpdong/imParalog.

## Table Legend

**Table S1.** Identification of T cell enhancing paralog pairs using the pSSEA method in six cancer cell CRISPR screen data.

**Table S2.** Experimental data of paralog pairs selected for validation in this study, including guide sequence, FACS data and statistical analysis of results.

**Table S3.** Construction of XGBoost classifier for predicting T cell enhancing paralog pairs.

## Supplemental Figure Legends

**Figure S1.**
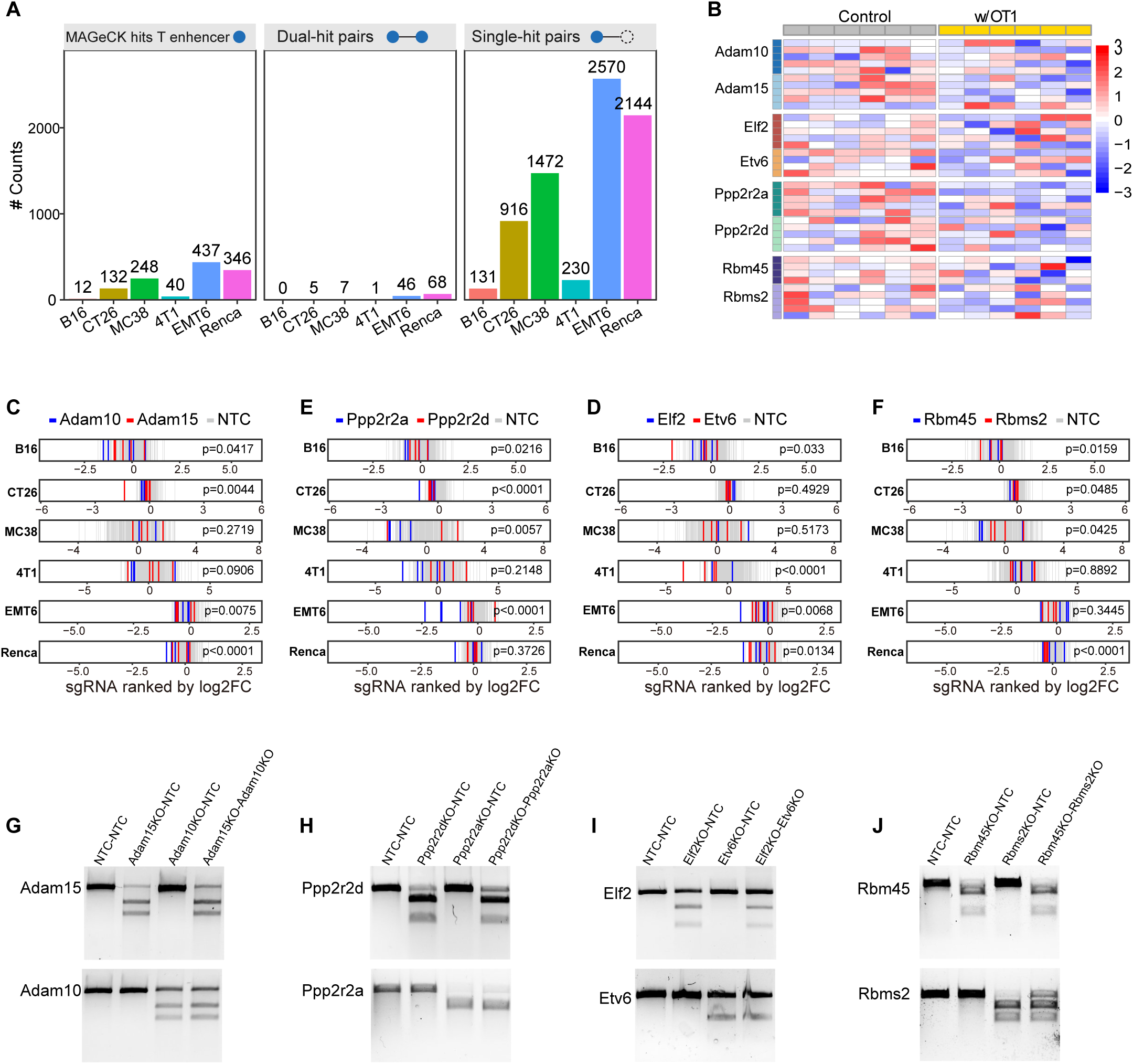
Validation of synergistic paralog pairs identified using pSSEA. (A) Selection of T cell enhancing paralog pairs using MAGeCK hits. Left panel: identification of cancer-intrinsic genes that enhance T cell killing using CRISPR screen data; Middle panel: Selection of dual-hit paralog pairs where both genes are identified as T cell enhancing hits; Right panel: Selection of single-hit paralog pairs where either gene is identified as a T cell enhancing hit. (B) Heatmap of B16F10 melanoma cell CRISPR-seq sgRNA profile from GSE149933 for selected paralog pairs. (C-F) Distribution of sgRNA ranked by log2FC for visualizing selected paralog pairs *Adam15/Adam10* (C), *Ppp2r2d/Ppp2r2a*(D), *Elf2/Etv6* (E) and *Rbm45/Rbms2* (F) in CRISPR-seq of six cell lines. The log2 fold change was measured by DESeq2 between with and without OT1 T cells co-cultured groups. (G-J) T7E1 assays were used to test the cutting efficiency of dual gene knockouts for *Adam15/Adam10* (G), *Ppp2r2d/Ppp2r2a*(H), *Elf2/Etv6* (I) and *Rbm45/Rbms2* (J).

**Figure S2.**
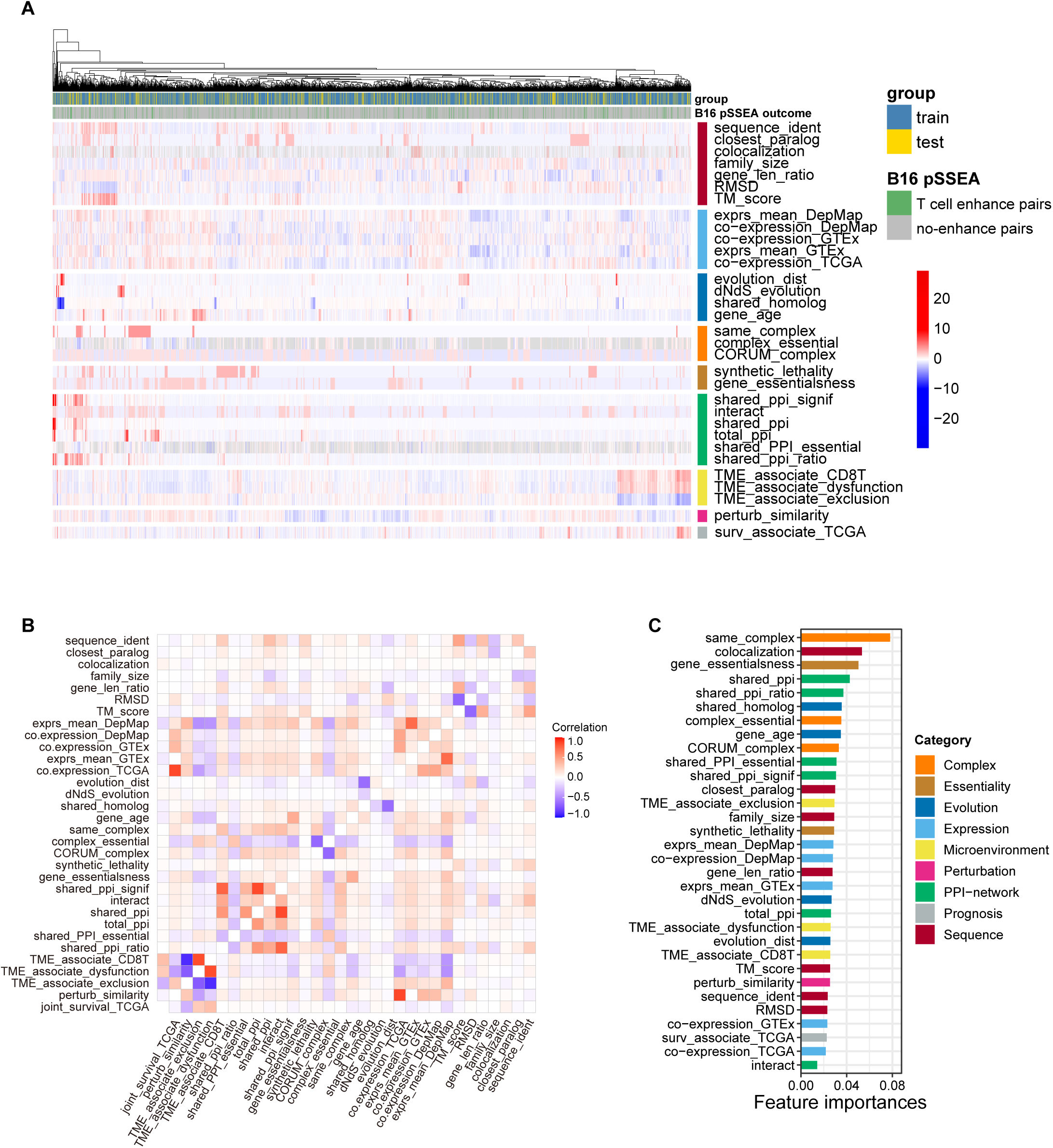
Data and feature characteristics for developing XGBoost classifier for predicting paralog pairs that synergistic enhance T cell function. (A) Heatmap of the training and the internal testing sets including all 3204 paralog pairs. (B) Correlation plot of 32 features used in the XGBoost classifier. The color represents the correlation between two features in 3204 paralog pairs. (C) Characterization of XGBoost classifier feature importance using the training set.

**Figure S3.**
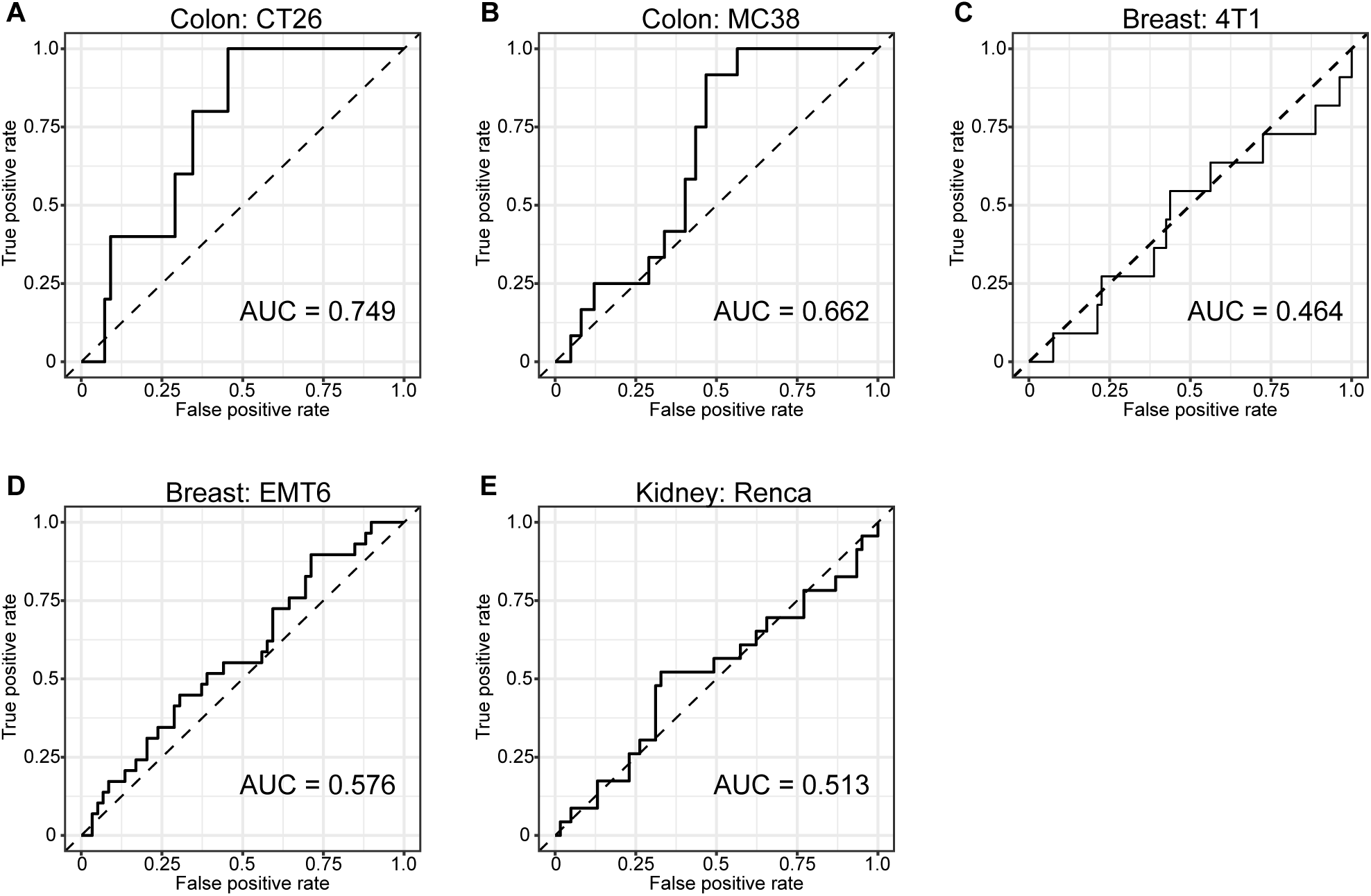
Evaluation of classifier performance on additional cancer cell lines. ROC curves showing the performance of our classifier in distinguishing paralog pairs out of B16F10 melanoma data in CT26 (A), MC38 (B), 4T1 (C), EMT6 (D), and Renca (E) cancer cell lines. Area under curve (AUC) was used for measuring the average precision of prediction.

**Figure S4.**
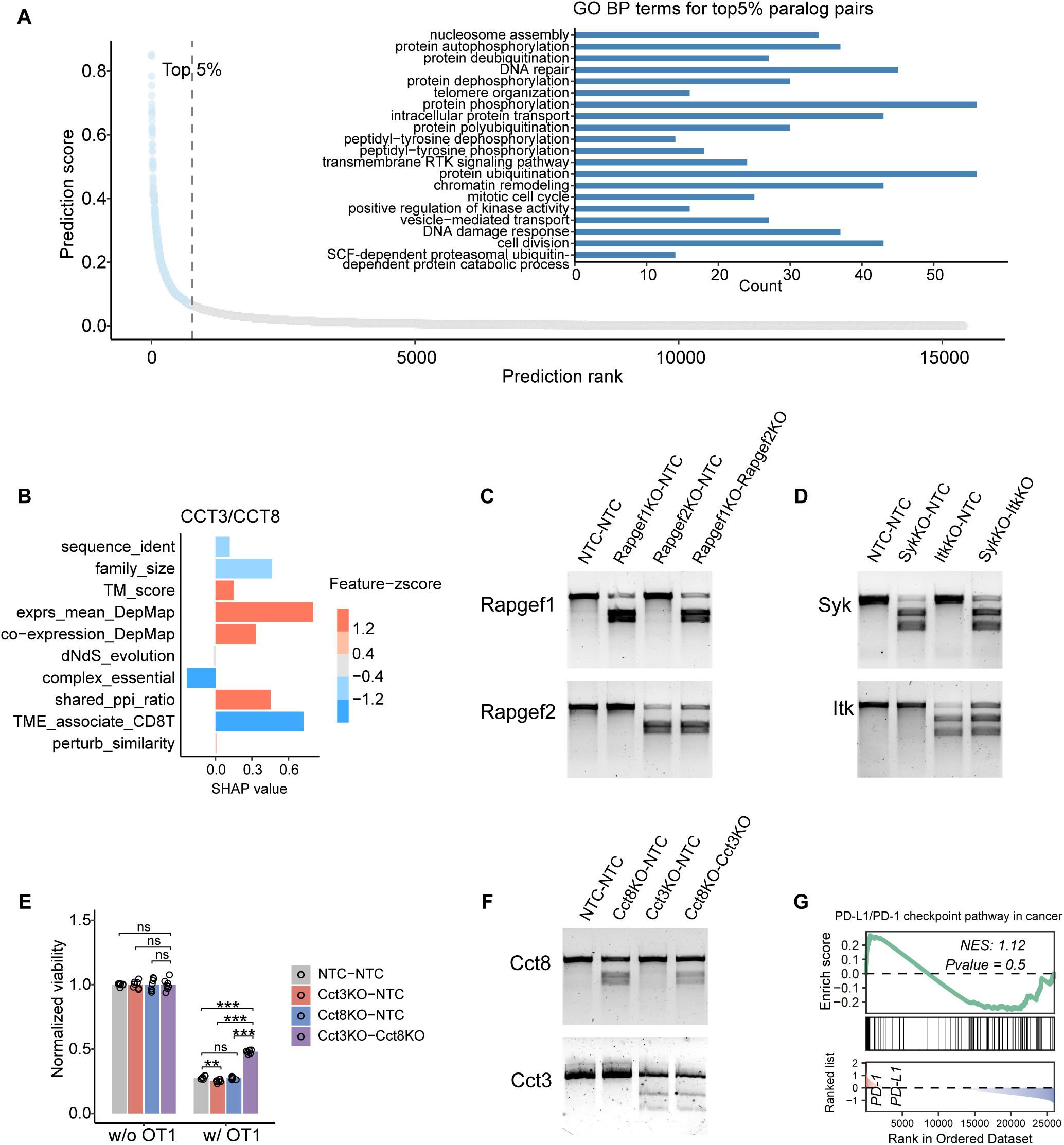
Test performance of top 3 predicted paralogs from XGBoost classifier. (A) Biological process pathways enriched in the top 5% of paralog pair genes based on prediction scores using clusterProfiler. (B) SHAP profile for *CCT3/CCT8*. (C, D) T7E1 assays were used to test the cutting efficiency of dual gene knockouts for *Rapgef1/Rapgef2* (C), and *Syk/Itk* (D). (E) Bar chart of mean viability of *Cct3/Cct8* knock-out treated B16F10 melanoma cells following culture with or without OT-1 T cells. The p values are from two-sided heteroscedastic t tests. (F) T7E1 assays were used to test the cutting efficiency of dual gene knockouts for *Cct3/Cct8*. (G) GSEA analysis between patients with low expression of *CCT3/CCT8* against PD-L1/PD-1 signaling pathway. The GSEA was performed using the differential expression output between patients low (<25%) in both *CCT3/CCT8* versus all remaining patients.

## References

1. Sharma, P., Hu-Lieskovan, S., Wargo, J.A., and Ribas, A. (2017). Primary, adaptive, and acquired resistance to cancer immunotherapy. Cell 168, 707–723.

2. Wang, G., Chow, R.D., Zhu, L., Bai, Z., Ye, L., Zhang, F., Renauer, P.A., Dong, M.B., Dai, X., and Zhang, X. (2020). CRISPR-GEMM pooled mutagenic screening identifies KMT2D as a major modulator of immune checkpoint blockade. Cancer discovery 10, 1912–1933.

3. Madani Tonekaboni, S.A., Soltan Ghoraie, L., Manem, V.S.K., and Haibe-Kains, B. (2018). Predictive approaches for drug combination discovery in cancer. Briefings in bioinformatics 19, 263–276.

4. Magen, A., Sahu, A.D., Lee, J.S., Sharmin, M., Lugo, A., Gutkind, J.S., Schäffer, A.A., Ruppin, E., and Hannenhalli, S. (2019). Beyond synthetic lethality: charting the landscape of pairwise gene expression states associated with survival in cancer. Cell reports 28, 938–948. e936.

5. Parrish, P.C., Thomas, J.D., Gabel, A.M., Kamlapurkar, S., Bradley, R.K., and Berger, A.H. (2021). Discovery of synthetic lethal and tumor suppressor paralog pairs in the human genome. Cell reports 36.

6. De Kegel, B., Quinn, N., Thompson, N.A., Adams, D.J., and Ryan, C.J. (2021). Comprehensive prediction of robust synthetic lethality between paralog pairs in cancer cell lines. Cell Systems 12, 1144–1159. e1146.

7. Park, J.S., Gazzaniga, F.S., Wu, M., Luthens, A.K., Gillis, J., Zheng, W., LaFleur, M.W., Johnson, S.B., Morad, G., and Park, E.M. (2023). Targeting PD-L2–RGMb overcomes microbiome-related immunotherapy resistance. Nature, 1–9.

8. Moreno-Lama, L., Galindo-Campos, M.A., Martínez, C., Comerma, L., Vazquez, I., Vernet-Tomas, M., Ampurdanés, C., Lutfi, N., Martin-Caballero, J., and Dantzer, F. (2020). Coordinated signals from PARP-1 and PARP-2 are required to establish a proper T cell immune response to breast tumors in mice. Oncogene 39, 2835–2843.

9. Park, J.J., Codina, A., Ye, L., Lam, S., Guo, J., Clark, P., Zhou, X., Peng, L., and Chen, S. (2022). Double knockout CRISPR screen for cancer resistance to T cell cytotoxicity. Journal of Hematology & Oncology 15, 1–5.

10. Zhao, D., Badur, M.G., Luebeck, J., Magaña, J.H., Birmingham, A., Sasik, R., Ahn, C.S., Ideker, T., Metallo, C.M., and Mali, P. (2018). Combinatorial CRISPR-Cas9 metabolic screens reveal critical redox control points dependent on the KEAP1-NRF2 regulatory axis. Molecular cell 69, 699–708. e697.

11. Tang, S., Wu, X., Liu, J., Zhang, Q., Wang, X., Shao, S., Gokbag, B., Fan, K., Liu, X., and Li, F. (2022). Generation of dual-gRNA library for combinatorial CRISPR screening of synthetic lethal gene pairs. STAR protocols 3, 101556.

12. Ito, T., Young, M.J., Li, R., Jain, S., Wernitznig, A., Krill-Burger, J.M., Lemke, C.T., Monducci, D., Rodriguez, D.J., and Chang, L. (2021). Paralog knockout profiling identifies DUSP4 and DUSP6 as a digenic dependence in MAPK pathway-driven cancers. Nature genetics 53, 1664–1672.

13. Lawson, K.A., Sousa, C.M., Zhang, X., Kim, E., Akthar, R., Caumanns, J.J., Yao, Y., Mikolajewicz, N., Ross, C., and Brown, K.R. (2020). Functional genomic landscape of cancer-intrinsic evasion of killing by T cells. Nature 586, 120–126.

14. Hänzelmann, S., Castelo, R., and Guinney, J. (2013). GSVA: gene set variation analysis for microarray and RNA-seq data. BMC bioinformatics 14, 1–15.

15. Edelman, E., Porrello, A., Guinney, J., Balakumaran, B., Bild, A., Febbo, P.G., and Mukherjee, S. (2006). Analysis of sample set enrichment scores: assaying the enrichment of sets of genes for individual samples in genome-wide expression profiles. Bioinformatics 22, e108–e116.

16. Aguirre-Ducler, A., Gianino, N., Villarroel-Espindola, F., Desai, S., Tang, D., Zhao, H., Syrigos, K., Trepicchio, W.L., Kannan, K., and Gregory, R.C. (2022). Tumor cell SYK expression modulates the tumor immune microenvironment composition in human cancer via TNF-α dependent signaling. Journal for ImmunoTherapy of Cancer 10, e005113.

17. Zhao, M., Li, L., Kiernan, C.H., Castro Eiro, M.D., Dammeijer, F., van Meurs, M., Brouwers-Haspels, I., Wilmsen, M.E., Grashof, D.G., and van de Werken, H.J. (2023). Overcoming immune checkpoint blockade resistance in solid tumors with intermittent ITK inhibition. Scientific Reports 13, 15678.

18. Thakur, R., Xu, M., Thornock, A., Sowards, H., Long, E., Rheling, T., Funderburk, K., Yin, J., Hennessey, R., and Chari, R. (2023). Integrative analysis of 3D chromatin organization at GWAS loci identifies RAPGEF1 as a melanoma susceptibility gene. Cancer Research 83, 5239–5239.

19. Vilella, A.J., Severin, J., Ureta-Vidal, A., Heng, L., Durbin, R., and Birney, E. (2009). EnsemblCompara GeneTrees: Complete, duplication-aware phylogenetic trees in vertebrates. Genome research 19, 327–335.

20. Boratyn, G.M., Camacho, C., Cooper, P.S., Coulouris, G., Fong, A., Ma, N., Madden, T.L., Matten, W.T., McGinnis, S.D., and Merezhuk, Y. (2013). BLAST: a more efficient report with usability improvements. Nucleic acids research 41, W29–W33.

21. Varadi, M., Anyango, S., Deshpande, M., Nair, S., Natassia, C., Yordanova, G., Yuan, D., Stroe, O., Wood, G., and Laydon, A. (2022). AlphaFold Protein Structure Database: massively expanding the structural coverage of protein-sequence space with high-accuracy models. Nucleic acids research 50, D439–D444.

22. Zhang, Y., and Skolnick, J. (2004). Scoring function for automated assessment of protein structure template quality. Proteins: Structure, Function, and Bioinformatics 57, 702–710.

23. Tsitsiridis, G., Steinkamp, R., Giurgiu, M., Brauner, B., Fobo, G., Frishman, G., Montrone, C., and Ruepp, A. (2023). CORUM: the comprehensive resource of mammalian protein complexes–2022. Nucleic acids research 51, D539–D545.

24. Capra, J.A., Williams, A.G., and Pollard, K.S. (2012). ProteinHistorian: tools for the comparative analysis of eukaryote protein origin. PLoS computational biology 8, e1002567.

25. Yang, Z. (2007). PAML 4: phylogenetic analysis by maximum likelihood. Molecular biology and evolution 24, 1586–1591.

26. Harikrishnan, S.L., Pucholt, P., and Berlin, S. (2015). Sequence and gene expression evolution of paralogous genes in willows. Scientific reports 5, 18662.

27. Tabach, Y., Golan, T., Hernández-Hernández, A., Messer, A.R., Fukuda, T., Kouznetsova, A., Liu, J.G., Lilienthal, I., Levy, C., and Ruvkun, G. (2013). Human disease locus discovery and mapping to molecular pathways through phylogenetic profiling. Molecular systems biology 9, 692.

28. Tabach, Y., Billi, A.C., Hayes, G.D., Newman, M.A., Zuk, O., Gabel, H., Kamath, R., Yacoby, K., Chapman, B., and Garcia, S.M. (2013). Identification of small RNA pathway genes using patterns of phylogenetic conservation and divergence. Nature 493, 694–698.

29. Sherill-Rofe, D., Rahat, D., Findlay, S., Mellul, A., Guberman, I., Braun, M., Bloch, I., Lalezari, A., Samiei, A., and Sadreyev, R. (2019). Mapping global and local coevolution across 600 species to identify novel homologous recombination repair genes. Genome research 29, 439–448.

30. Gurumayum, S., Jiang, P., Hao, X., Campos, T.L., Young, N.D., Korhonen, P.K., Gasser, R.B., Bork, P., Zhao, X.-M., and He, L.-j. (2021). OGEE v3: Online GEne Essentiality database with increased coverage of organisms and human cell lines. Nucleic acids research 49, D998–D1003.

31. Dempster, J.M., Boyle, I., Vazquez, F., Root, D.E., Boehm, J.S., Hahn, W.C., Tsherniak, A., and McFarland, J.M. (2021). Chronos: a cell population dynamics model of CRISPR experiments that improves inference of gene fitness effects. Genome biology 22, 1–23.

32. Ghandi, M., Huang, F.W., Jané-Valbuena, J., Kryukov, G.V., Lo, C.C., McDonald III, E.R., Barretina, J., Gelfand, E.T., Bielski, C.M., and Li, H. (2019). Next-generation characterization of the cancer cell line encyclopedia. Nature 569, 503–508.

33. Lonsdale, J., Thomas, J., Salvatore, M., Phillips, R., Lo, E., Shad, S., Hasz, R., Walters, G., Garcia, F., and Young, N. (2013). The genotype-tissue expression (GTEx) project. Nature genetics 45, 580–585.

34. Weinstein, J.N., Collisson, E.A., Mills, G.B., Shaw, K.R., Ozenberger, B.A., Ellrott, K., Shmulevich, I., Sander, C., and Stuart, J.M. (2013). The cancer genome atlas pan-cancer analysis project. Nature genetics 45, 1113–1120.

35. Subramanian, A., Narayan, R., Corsello, S.M., Peck, D.D., Natoli, T.E., Lu, X., Gould, J., Davis, J.F., Tubelli, A.A., and Asiedu, J.K. (2017). A next generation connectivity map: L1000 platform and the first 1,000,000 profiles. Cell 171, 1437–1452. e1417.

36. Jiang, P., Gu, S., Pan, D., Fu, J., Sahu, A., Hu, X., Li, Z., Traugh, N., Bu, X., and Li, B. (2018). Signatures of T cell dysfunction and exclusion predict cancer immunotherapy response. Nature medicine 24, 1550–1558.

37. Sen, P.K. (1968). Estimates of the regression coefficient based on Kendall’s tau. Journal of the American statistical association 63, 1379–1389.

38. Mounir, M., Lucchetta, M., Silva, T.C., Olsen, C., Bontempi, G., Chen, X., Noushmehr, H., Colaprico, A., and Papaleo, E. (2019). New functionalities in the TCGAbiolinks package for the study and integration of cancer data from GDC and GTEx. PLoS computational biology 15, e1006701.

39. Chen, T., and Guestrin, C. (2016). XGBoost: A Scalable Tree Boosting System. Proceedings of the 22nd ACM SIGKDD International Conference on Knowledge Discovery and Data Mining. Association for Computing Machinery.

